# Analysis of protein levels and solubility in distinct brain regions reveals several elements of the protein homeostasis network that are impacted by aging

**DOI:** 10.1101/2024.02.28.582597

**Authors:** Cristen Molzahn, Erich Kuechler, Lorenz Nierves, Gary Cole, Jing Wang, Neil Cashman, Aly Karsan, Philipp F. Lange, Jörg Gsponer, Thibault Mayor

## Abstract

The onset of protein conformation diseases is inextricably linked to aging. During aging, cellular protein quality control declines which results in diminished protein homeostasis (proteostasis). In model organisms, such as *C. elegans* and killifish, proteostatic decline with age has been linked to the onset of aggregation of proteins in wild-type animals, observed through detergent-insoluble fractionation. Analysis of studies applying detergent-insoluble fractionation in mice revealed that the composition of detergent-insoluble proteins changes with age. However, these individual fractionation studies have generally been limited to small numbers of mice. Herein, we expand on our previous analysis by extending the experiments to a larger cohort of mice and to two brain regions implicated in neurodegenerative diseases, the cortex and hippocampus. These experiments unveil insights into alterations in the abundance and solubility of proteins involved in protein quality control and in inflammation. For example, ribosomal proteins and many chaperone proteins are downregulated with age. Consistent enrichment of subunits of the extracellular C1q complex was also observed in both brain regions alongside an increase in immunoglobulin signal indicating that markers of increased inflammation may also become insoluble during aging. More generally, insoluble proteins share features observed in datasets of impaired protein degradation indicating that the loss of activity of cellular protein degradation machinery may contribute to the specific aggregation of these proteins.

## Introduction

Declines in cellular protein quality control collectively give rise to the loss of protein homeostasis that is implicated in aging and neurodegenerative diseases (NDs). Loss of activity of chaperones, autophagy and the proteasome contribute to the accumulation of damaged, misfolded and ubiquitinated proteins within protein aggregates. Characterization of the composition of aggregates as well as age-related changes to protein quality control (PQC) is needed to better understand the mechanisms of age-related protein aggregation.

The age-related decline in proteostasis is hypothesized to promote protein conformation diseases. Loss of proteostasis and the formation of detergent-insoluble protein aggregates have been widely observed with age.^1^ Such detergent-insoluble aggregates also characterize NDs.^2^ Modulation of PQC components has been demonstrated to extend lifespan in model organisms. For example. reducing translation errors by introducing ribosome mutations that increase fidelity extends lifespan in yeast, worms and flies.^3^ Enhancing HSP70 levels through exogenous expression improves synapse function, memory and extends lifespan in mice.^4^ In flies, overexpression of proteasome subunits in neurons extends lifespan and enhances memory late in life.^5^ Additionally, canonical aging pathways frequently influence protein metabolism. The formation of aggregates in protein conformation diseases is highly cell-type and brain region specific.^6^

The decline of PQC machinery is heterogeneous, occurring in cell type and tissue-specific patterns. In a study of microarray data in healthy brain tissue, chaperones that were previously shown to influence Aβ aggregation were observed to be expressed at lower levels in regions impacted by Alzheimer’s disease (AD) and an increase in expression was observed for proteins previously found to co-aggregate with Aβ plaques and neurofibrillary tangles (NFTs).^7^ Aggregation-prone proteins effect neuron types selectively, remaining soluble in certain cells while aggregating in others.^8^ Specific neuronal vulnerability may be the site for onset of NDs, but aggregates can be transmitted to nearby cells in a prion-like seeding mechanism leading to reproducible patterns of aggregation.^9^ Regions of the cortex accumulate aggregates in several neurodegenerative diseases such as Creutzfeldt–Jakob disease, Lewy body disorders and Frontotemporal Lobar Degeneration.^6^ The hippocampus is affected by Aβ aggregates early in Alzheimer’s disease.^10^ Changes to protein levels and solubility can clearly predispose brain tissues to NDs in a region-specific manner. Reduced degradative capacity of the proteasome may contribute to the age-related increase in half-life of some proteins.^11^

We previously assessed by mass spectrometry (MS) the presence of triton-insoluble proteins in mouse brains and showed that a small subset of the proteome becomes more insoluble upon aging.^12^ These age-associated insoluble proteins tend to be more structured and depleted for intrinsic disordered. Proteins with a similar physicochemical “signature” are found to be insoluble in several other proteomics mouse studies of different ND models, but only when older mice are assessed.^12^ In contrast, proteins more insoluble in brain tissue of younger mice tend to be more disordered and associated with membraneless organelles (MLO; *i.e.,* cellular condensates) as compared to older mice. Therefore, while some MLO-associated proteins are generally more insoluble, we saw no evidence for an age-dependent accumulation of these proteins in the insoluble fraction. This finding is intriguing since it was previously suggested that changes in the liquid-like properties of MLOs could lead to the accumulation of aggregated proteins upon aging and in NDs.^13^ Instead, our work indicates that more hydrophobic proteins are enriched in the detergent-insoluble fraction upon aging. Nonetheless, our first analysis was limited to a relatively small cohort of older mice and did not consider the overall age-associated changes in protein abundances.

In this work, we examined protein solubility in a region-specific manner using a larger number of mice to better account for possible variability. We identified detergent-insoluble proteins from two brain regions, the hippocampus and cortex. The insoluble proteins share similar GO terms and protein features indicating commonly affected pathways and proteins. Intriguingly, elements of the complement system and immunoglobulin proteins were identified in the detergent-insoluble fraction of older mice. Furthermore, we found levels of ribosomal proteins and of numerous chaperone proteins in both analyzed regions were lower in older mice, which could explain why some proteins become more insoluble upon aging. Finally, many of the detergent-insoluble proteins overlap with prominent components of lipofuscin that may accumulate in tissues of older mice.

## Results

### Identification of proteins with reduced solubility in the hippocampus and cortex due to aging

In order to identify differences in protein solubility between brain regions, detergent-insolubility fractionation MS was used in the cortex and hippocampus. A large cohort of mice (n = 13 old, 100-weeks and n = 15 young, 15 weeks) was analyzed to identify effect sizes with greater confidence. Tissue samples were lysed in a 1% triton containing buffer and subjected to centrifugation to isolate a detergent-insoluble fraction. The content of these fractions (i.e., supernatant and pellet) was analyzed by data independent acquisition (DIA)-MS (Figure 1A). From both brain regions and fractions, 6,925 protein groups and 83,420 peptides were quantified. 6,847 and 6,822 protein groups were quantified in the hippocampus and cortex, respectively, with 65% quantified in > 90% of the samples (Table S1). The resulting data shows strong correlations between protein group intensities of biological replicates, with a few exceptions (Figure S1A-B). After removal of 4 samples with poor data quality, median R squared values within age and fraction groups ranged from 0.86-0.97. Principal component analysis (PCA) shows that the data was separated by fractions and brain regions, creating 4 clusters (Figure S1C). This finding suggests that the composition of the two fractions is influenced by brain region. Additionally, this analysis reveals differences in the data obtained from both fractions. The supernatant fraction data clusters strikingly more tightly as compared to the pellet fraction data, especially in the first dimension. This suggests that individual supernatant datasets are more similar to one another, while pellet fraction sample may have more variability. Comparing only the pellet fraction, the samples are still strongly clustered by brain regions (Figure 1B). Importantly, when PCA is performed on a single brain region and fraction, there is no clustering of the data by age (Figure S1 D-G), indicating that protein composition of each fraction is not drastically changing and that only a subset of proteins undergoes a solubility shift during aging.

**Figure 1.**
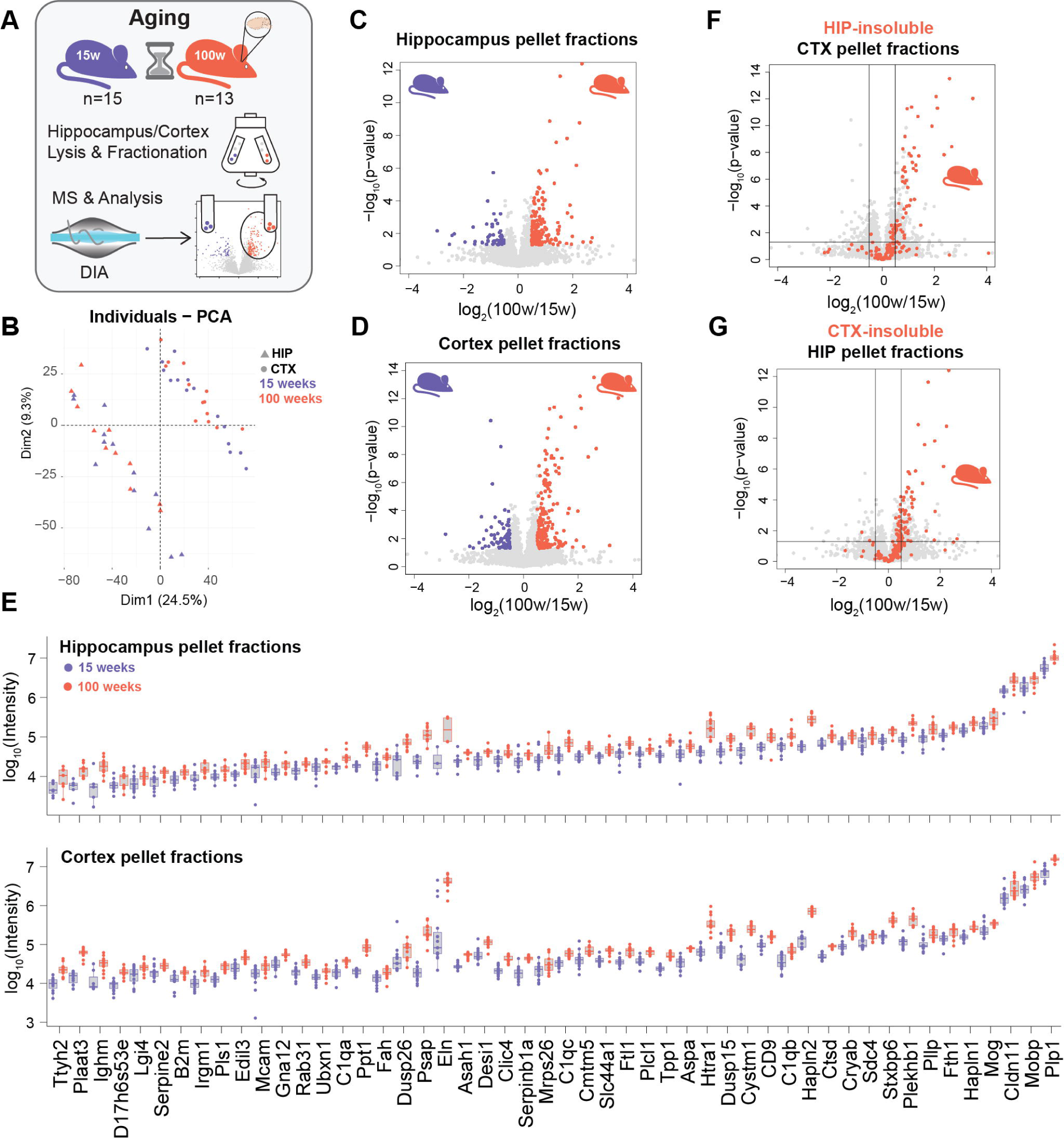
Changes in levels of detergent-insoluble protein occur in a subset of the observed proteome. A) Schematic representing experimental setup. The red mouse and data points denote results from older mice. B) PCA of protein group intensities measured in the pellet fraction of the hippocampus (HIP) and cortex (CTX), excluding 2 data sets with low correlation. Significantly altered proteins between mice ages 100- and 15-weeks-old identified in the C) hippocampus pellet fraction D) cortex pellet fraction. Significance was determined using a two-sample t-test assuming equal variance. The p-value cut off was set as 0.05 and the fold change at a log_2_ ratio of 0.5. E) Individual log-transformed protein group intensity values for each mouse plotted for each of the 50 common age-associated detergent-insoluble proteins analyzed in the hippocampus and cortex pellet fractions. Datapoints are divided by age group with the 100-week-old mice shown in red and the 15-week-old mice shown in purple. F) the 152 age-associated detergent-insoluble proteins in the hippocampus mapped onto their ratios in the cortex pellet fraction and G) The 179 detergent-insoluble proteins in the cortex mapped onto their corresponding ratios in the hippocampus pellet fraction.

To identify a subset of proteins that experience solubility changes associated with aging, the fractions were compared between young and old mice using a t-test. This analysis revealed a group of 152 and 197 proteins that were more insoluble in old mice in the hippocampus and the cortex, respectively (Figure 1C-D). We also identified 57 and 82 proteins that were enriched in the insoluble fraction of the hippocampus and cortex in young mice. These results are similar to what we observed using a smaller number of mice in an independent mouse cohort.^12^ This recapitulation indicates that, perhaps surprisingly, using a larger number of mice (13-15 vs 3-4), did not allow the identification of a higher number of detergent-insoluble proteins affected by aging, possibly due to the high variability in the pellet fractions, and the small number of proteins affected by aging. Importantly, the aspartoacylase (ASPA) and the alpha-crystallin B chain (CRYAB) small heat shock protein, which were previously validated using a filter trap assay (FTA), were also found to be more detergent-insoluble upon aging in this second analysis. In the supernatant, which also represents a whole cell lysate analysis, similarly small changes in abundance were observed with 90 proteins increasing in the hippocampus and 58 protein groups significantly depleted (Figure S2A). In the cortex, 99 proteins increase in abundance and 72 decrease (Figure S2B). These results confirm that there are no striking shifts in protein expression levels upon aging in brain tissues, consistent with previous work and our published results.^12, 14, 15^

We next sought to identify regional differences between the cortex and hippocampus by comparing the composition of the detergent-insoluble fractions significantly affected by aging. A total of 50 proteins are significantly increased upon aging in both pellet fractions (Figure S2C). Therefore, less than a third of proteins that become more insoluble in one tissue are also affected in the second analyzed brain region upon aging. To better evaluate the abundance of these proteins in the pellet fraction, we plotted the log transformed label-free intensities of the 50 insoluble proteins identified in both tissues (Figure 1E). These proteins often share similar intensities among the different mice and across brain regions. In both regions, the three proteins with prominently higher intensities in the pellet fractions (regardless of the age) are PLP1 (myelin proteolipid protein), MOBP (myelin-associated oligodendrocyte basic protein) and CLDN11 (claudin 11). This enrichment is reminiscent of the composition of lipofuscin the formation of which, in microglia, may be driven by the engulfment of myelin.^16^ The remaining 102 and 129 proteins appear specific to the hippocampus and cortex, respectively (Figure S2C). Nonetheless, a substantial portion of proteins enriched with age in the detergent-insoluble fraction of one brain region trends towards enrichment in the detergent-insoluble fraction with age in the other (Figure 1F-G). These results indicate that many age-dependent detergent-insoluble proteins become prominently enriched in one tissue while also being affected in the other tissue, but typically below our set threshold.

To determine whether these proteins are associated with different cellular functions we performed GO analysis of each group. The group of 152 proteins enriched in the detergent-insoluble fraction of the hippocampus contained a large share of lysosomal proteins, extracellular proteins and proteins involved in autophagosome assembly (Figure 2A). Among the 179 proteins enriched in the detergent-insoluble fraction from the cortex, enrichment was observed for lysosomal and extracellular proteins indicating that similar proteins are affected across brain regions. This group of lysosomal proteins includes the protein ATP6V0c, which when knocked down in neuroblastoma cells leads to lysosomal deficits, disrupted lysosomal pH and the accumulation of APP and α-syn.^17^ Disruption of lysosomal pH in aged brain tissue may also explain the presence of lysosomal proteins in the detergent insoluble fraction. Alkaline lysosomes have been documented to be more permeable leading to cytosolic localization of lysosomal proteins such as cathepsins.^18^ Cathepsins are proteases that are typically active in the low pH environment of the lysosome. Several cathepsin proteins were identified in the detergent-insoluble fractions from both the hippocampus and cortex. Collectively these findings suggest that changes in solubility of lysosomal proteins may reflect alterations in lysosomal function leading to reduced degradation. The loss of solubility of proteins from these compartments may contribute to the decline in these organelles that has previously been observed with age and may overlap with the well-characterized accumulation of lipofuscin. Lipofuscin is a detergent-insoluble inclusion consisting mainly of protein (30-70%) that forms due to defective lysosomal degradation.^19, 20^ Proteins that are specific to the hippocampus (102 proteins) are enriched for lysosomal proteins and those that are specific to the cortex (129 proteins) are enriched for mitochondrial proteins. GO analysis of the 50 common proteins reveals an enrichment for lysosomal proteins, hydrolases, extracellular proteins and members of the C1 complex (Figure 2A). In fact, 26% of the hippocampus proteins (40/152) and 20% of the cortex proteins (35/179) are annotated to localize extracellularly. The enrichment of extracellular proteins is surprising, since we initially aimed at identifying features associated with proteins that misfold and aggregate in the cell. Nonetheless, without carefully evaluating where these proteins aggregate, it remains unclear where these detergent-insoluble proteins are located.

**Figure 2.**
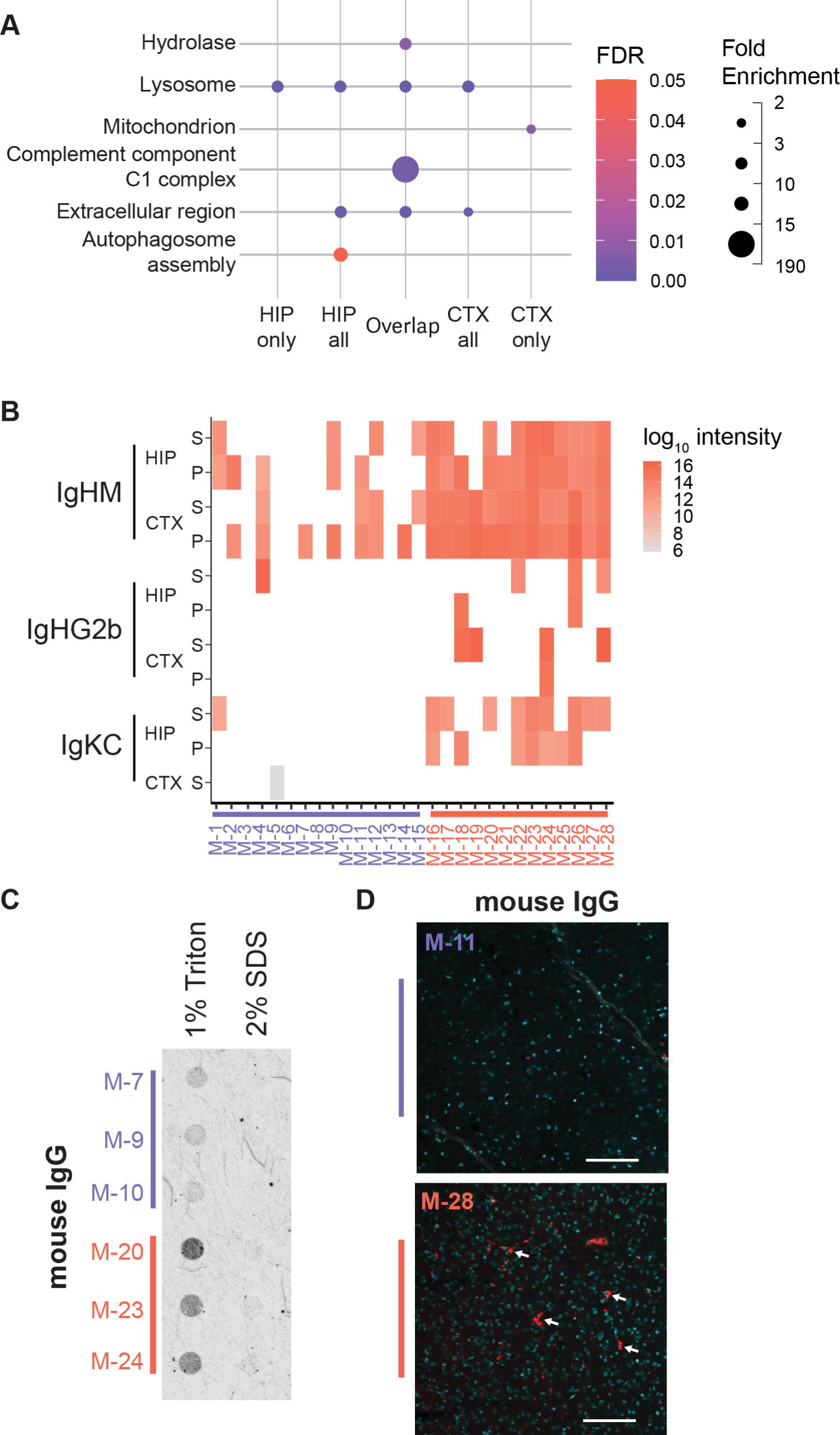
Immunoglobulins increase in abundance and become insoluble in old mice. A) Gene ontology (GO) analysis showing enrichment for age-associated detergent-insoluble proteins specific to the hippocampus (HIP; 102), specific to the cortex (CTX; 129) and shared between both regions (50). FDR values are Benjamini-Hochberg corrected p-values. B) Log_2_ transformed label-free intensities for each of the immunoglobulins identified by MS for each mouse where they were detected. Mice 1-15 (M-1 to M-15) are 15-weeks-old and mice 16-28 (M-16- to M-28) are 100-weeks-old. C) Filter trap assay of cortex-derived lysates of the indicated mice with immunodetection by goat anti-mouse IgG (1:10,000). D) Zoomed images, also shown in Figure S4, of the mouse cortex stained with goat anti-mouse IgG (1:200) and DAPI taken with a 10x objective from a young mouse (M-11) and an old mouse (M-28). Scale bar is 100 µm.

### Increased C1 and immunoglobulin signals are consistently detected in aged mice

We next sought to better understand the mechanism of C1q insolubility by better characterizing a phenomenon observed in our aging tissue. The C1 complex consists of C1r2, C1s2 and C1q. The C1q complex has three subunits, C1qa, C1qb and C1qc all of which are identified in the detergent-insoluble fraction in both the cortex and hippocampus. High levels of these proteins are also observed in both tissues (Figure S2A-B). The complement pathway of the innate immune system is activated by the binding of the C1q complex to antibodies in complex with antigens or to proteins such as Aβ and hyperphosphorylated Tau.^21^ Alterations in C1q levels with age has been implicated in the removal of synapses during aging. In aged mouse brain tissue, C1q has been observed to increase in abundance and colocalize with synapses. Disruption of this effect by knocking out C1q in mice improves memory in aged animals.^22^ Interestingly, proteins that are annotated as localized to the postsynaptic-density or to synapse trend toward lower expression in older mice hippocampus (Figure S2D-E). Increased expression of the C1q complex in the older mice may contribute to the loss of synapses observed in aged mice.

Along with the observed increase of complement proteins, levels of several antibodies become higher upon aging in both tissues (Figure 2B). Peptides matched three circulating antibody proteins: the immunoglobulin heavy constant mu (IGHM), the immunoglobulin heavy constant gamma 2 (IGHG2) and the immunoglobulin kappa constant chain (IGKC). Four of the young mice did not have any antibodies detected, whereas all of the old mice had at least one immunoglobulin polypeptide detected (Figure 2B). Notably, IGHM is prominently detected in both the hippocampus and cortex of the older mice, especially in the pellet fractions. These antibody proteins may result from previously observed disruption of the blood brain barrier with age. However, they may also be secreted by brain immune cells.^23^ IGKC has been previously observed to increase in abundance with age across mouse tissues.^15^ It was also recently shown that the IGHM constant domain is widely transcribed in mice brain, potentially from a cryptic start transcription site.^24^ The mu domain is the first of the 8 different mouse constant domains located downstream of the V_H_, D_H_, and J_H_ DNA regions that recombine during the differentiation of the B Cells. Expression of a neuronal IgM was proposed to have a role in cell adhesion or complement system activation.^24^ Notably, we detected peptides mapping both the N and C-terminal regions of the mu domain (Figure S3), indicating the whole domain translation. While these antibody proteins are detected by MS in both the supernatant and pellet fractions, FTA shows that IgGs are readily retained on cellulose acetate membranes in older mice (Figure 2C), suggesting that they are part of large, triton-insoluble complexes. Additionally, immunofluorescence (IF) staining with the anti-mouse secondary antibody reveals much more background signal in the aged mouse cortex as compared to the young mouse cortex (Figure 2D, Figure S4). Notably, the antibody signal forms bright foci that are similar in size to nuclei. Together these observations suggest that antibodies are increasing in the brain with age and may be colocalized with large, triton-insoluble foci. As IgM are well characterized C1q ligands^25^, their low solubility may contribute to the reduced solubility observed in the C1q complex components. These foci may also reflect the oligomerization of immunoglobulins that occurs when they bind to an antigen which enhances the binding of C1q.^26^

### Feature analysis of detergent-insoluble proteins

We next sought to analyze the features associated with detergent-insoluble proteins in both tissues. We previously demonstrated that, while different proteins were identified in the insoluble fractions obtained with distinct fractionation conditions or from different ND studies using old mouse models, these proteins share common features.^12^ Using the same 14 features that were identified in our previous study, the two populations of detergent-insoluble proteins derived from either the hippocampus or the cortex are markedly separated based on age in a PCA diagram with 72.3% of the variation captured in dimension 1 (Figure 3A).^12^ The insoluble proteins enriched in the old mice are correlated with features such as β-sheet content, percentage of sequence in Pfam domains and hydrophobicity. Similarly, detergent-insoluble proteins in aged mice are significantly depleted for intrinsic disorder and have more of their sequence in Pfam domains when compared to the whole mouse proteome (Figure 3B). Furthermore, proteins that become detergent-insoluble with age in both regions are depleted for random coils. In the hippocampus, detergent-insoluble proteins from old mice are enriched for β-sheet content and hydrophobicity. Detergent-insoluble proteins from the aged cortex show similar trends, however, these trends are not significant. Proteins enriched in the detergent-insoluble fraction from younger mice from both tissues display the opposite trends in comparison to proteins insoluble in older mice (Figure 3B). When considering features associated with solubility, the greatest effects were observed in the detergent-insoluble proteins from the young and old hippocampus. Similarly, insoluble proteins from the hippocampus of old mice are enriched for aggregation prone regions. While result trends are similar the differences are not significant in the cortex data. Overall, these results suggest that while the composition of proteins that become more enriched upon aging in the detergent-insoluble fractions in both tissues is largely distinct, they share very similar structural features.

**Figure 3.**
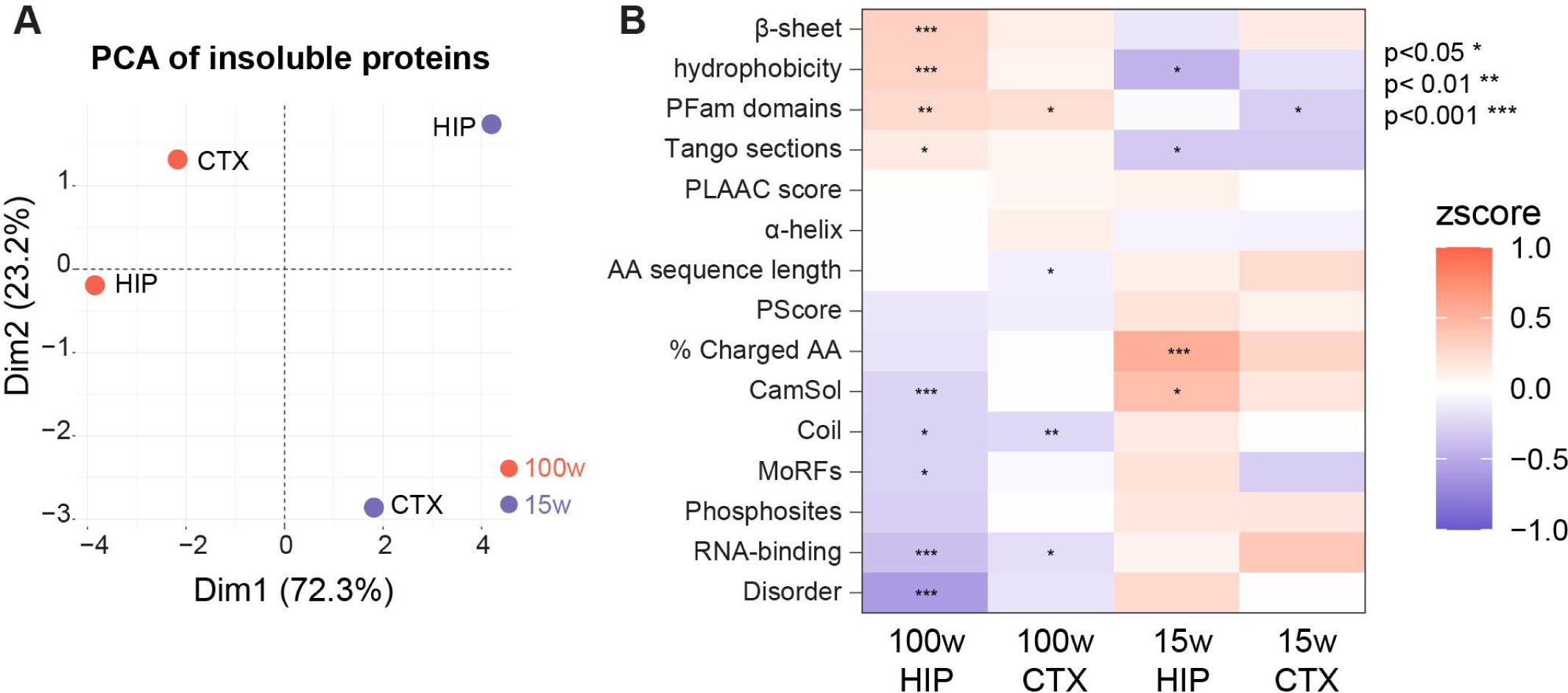
Age-associated detergent-insoluble proteins share common features associated with secondary structure and solubility. A) PCA of the feature means calculated for the groups of detergent-insoluble proteins based on ages and tissues (HIP: hippocampus; CTX: cortex). B) Individual feature distributions are shown for detergent insoluble proteins in 100-week-old (red) and 15-week-old (purple) mice. The number of proteins in each group is displayed below each violin plot. P-values (Wilcocxon test corrected using Benjamini-Hochberg method) are obtained from tests against the proteome distribution (grey).

Interestingly, many age-associated insoluble proteins tend to be more abundant in the respective tissues (Figure S5A, B). However, not all proteins that are more abundant in these two tissues are enriched in the triton-insoluble pellets (Figure 4A-B). To determine the role these features might play in predisposing a protein to aggregation, we assessed the features associated with proteins that increase in abundance in the supernatant alone and those that increase in both fractions. Proteins that increase in abundance in both the pellet and supernatant fractions share similar features to detergent-insoluble proteins in general. This group of proteins is more abundant as compared to the proteome and all soluble proteins that increase in expression (Figure 4B). They are depleted for intrinsic disorder while also showing a trend towards more β-sheet content and enrichment for Pfam domains (Figure 4C-E). By contrast, proteins that only increase in abundance in the supernatant are more similar to the whole proteome and do not show significant trends in the features assessed (Figure 4B-E). This data suggests that the features we have identified may predispose a protein to become insoluble upon increased expression.

**Figure 4.**
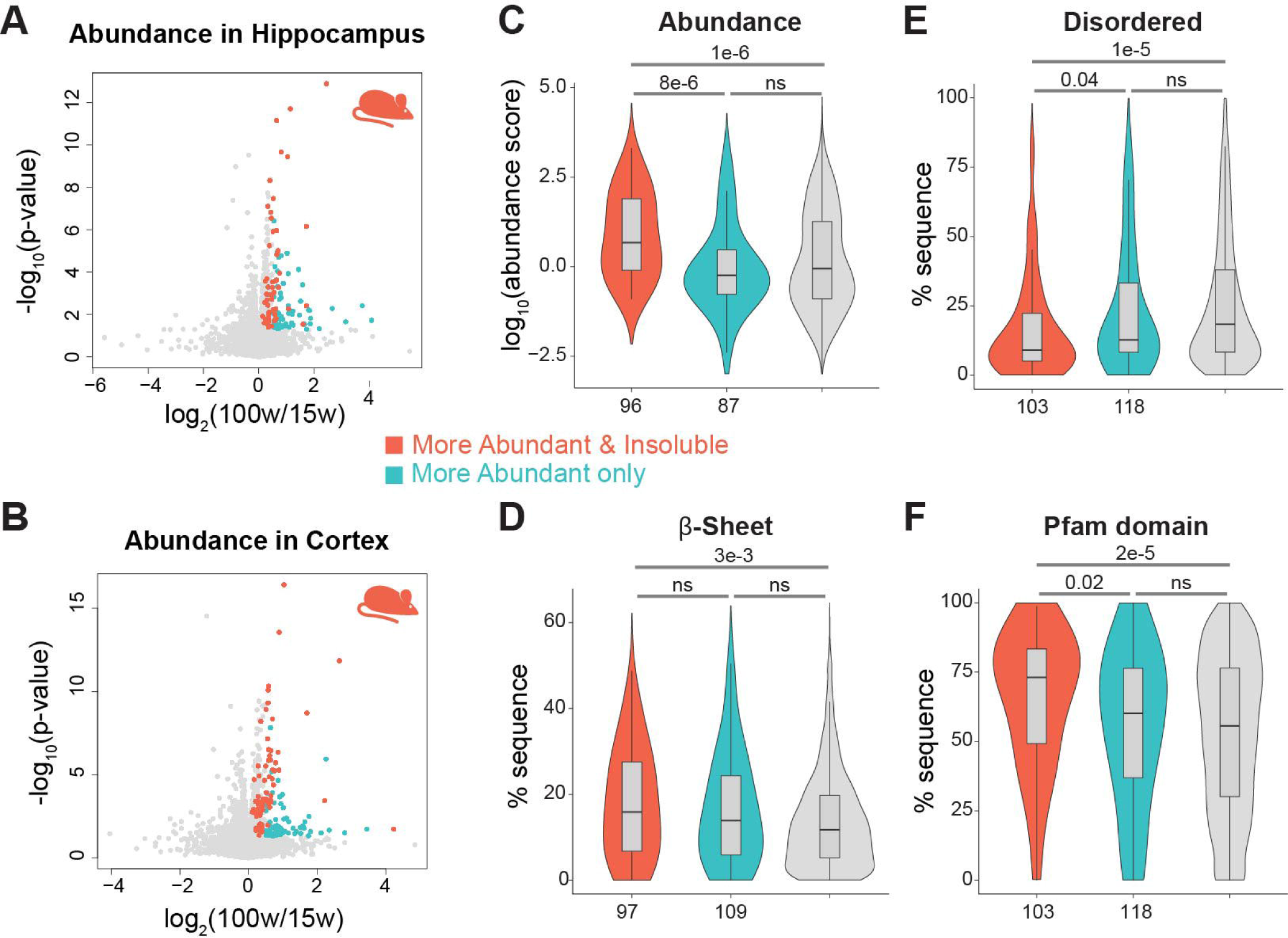
A subset of more abundant proteins also become more detergent-insoluble upon aging. A-B) volcano plot representing the age-associated changes in protein abundance in the hippocampus (A) and cortex (B). C-F) Proteins significantly more abundant but not more insoluble upon aging are shown in blue, while proteins also significantly enriched in the pellet fractions are shown in red. Feature distributions of indicated proteins that are more abundant upon aging in the hippocampus and cortex are shown for abundance (C), percent of the sequence predicted to form β-sheet (D), percent of the sequence predicted to be disordered (E), percent sequence within Pfam domains (F). P-values obtained from Hochberg adjusted Wilcoxon test.

### PQC components show altered expression with age

To gain more insights into protein quality control across the tissue samples and fractions, specific GO terms were compared. This analysis allows for the identification of alterations and potential changes in regulation of PQC pathways. Protein synthesis is affected by aging across organisms.^27, 28^ Strikingly, levels of ribosome proteins decrease upon aging in the hippocampus and cortex (Figure 5A). In cortex pellet fraction samples, ribosome levels also increase.

**Figure 5.**
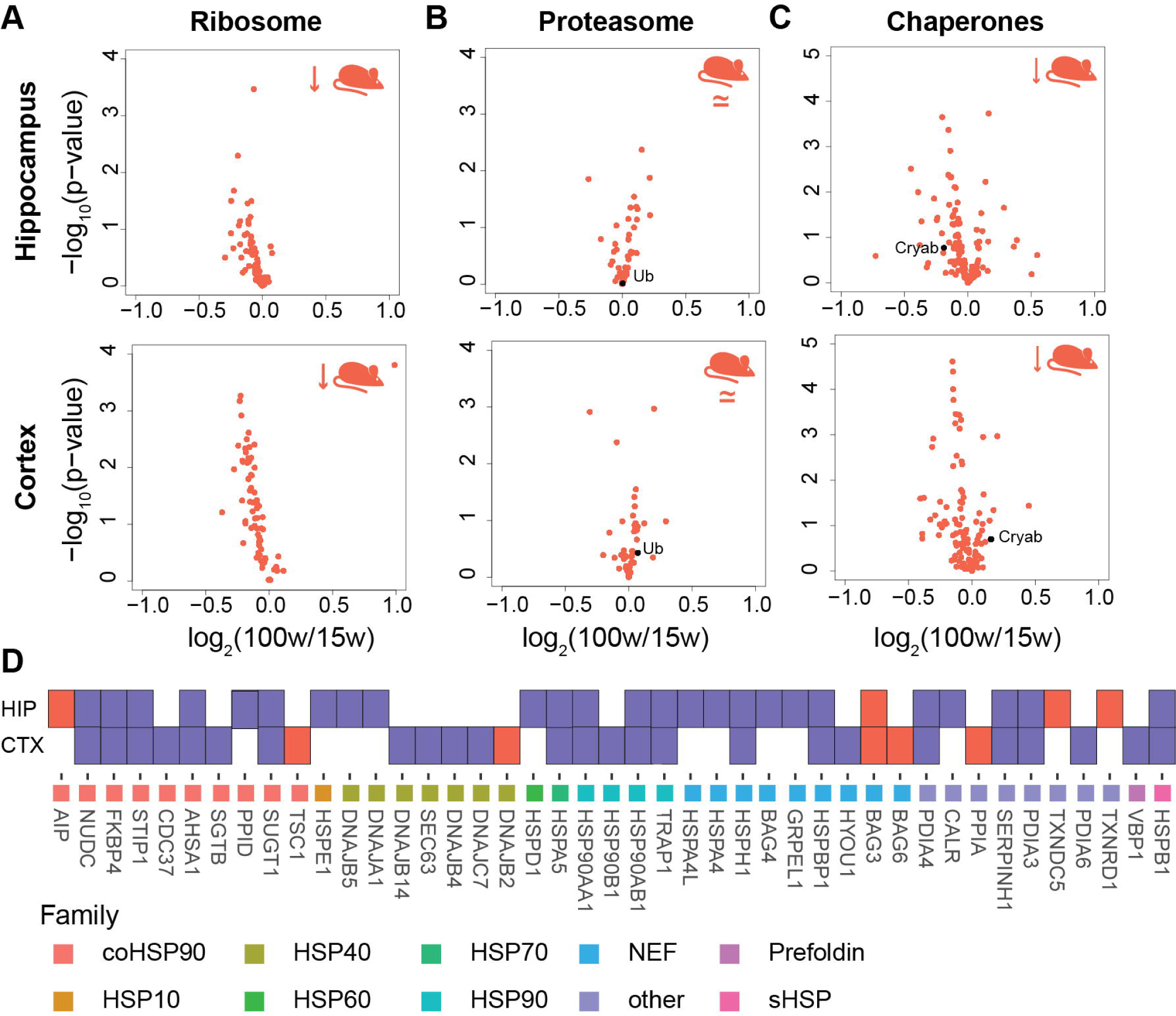
Alterations in the expression of PQC components. A-C) Proteins annotated with the indicated GO terms (A: ribosomal proteins, B: proteasomal subunits and ubiquitin (Ub), C: chaperones) are plotted according to their log_2_(fold-change) and t-test p-value to highlight changes in their expression levels. The corresponding Volcano plots for all proteins are shown in Figure S2. D) List of chaperone significantly (p<0.05) altered regardless of fold change (depleted: purple; enriched: red) in hippocampus (HIP) or cortex (CTX).

However, no similar trend was observed in the hippocampus (Figure S6A). The overall reduction of ribosome levels in the supernatant suggests that protein synthesis is potentially impaired, possibly to a higher extent in the aging cortex. In contrast, levels of proteasomal subunits do not show strong trends (Figure 5B). Curiously, these subunits show opposite trends in the pellet fractions, where they are either enriched or depleted in the hippocampus and cortex of older mice (Figure S6B). There is an accumulation of ubiquitin in the detergent-insoluble fraction in the cortex of old mice indicating that ubiquitinated proteins may be accumulating in the insoluble fraction (Figure S6B).

Overall levels of chaperone proteins of various classes were decreased with some exceptions of a few specific proteins such as the BAG3 co-chaperone that is present at higher levels in both tissues (Figure 5C-D). Notably, BAG3 is expressed at higher levels in an aging cell model and older mice, where it enhances autophagy.^29^ The depletion was observed across most chaperone classes. It includes several Hsp90s and Hsp90 co-chaperones, as well as the HSPH1 and HSPBP1 nucleotide exchange factor (NEF) that are expressed at slightly lower levels in both tissues (Figure 5C-D). The slight age-associated reduction of these chaperone proteins could lead to a lower folding capacity and the accumulation of their clients in the detergent-insoluble fraction. In addition, the small heat shock protein HSPB1 (a.k.a. Hsp27) is present at lower levels in older mice (17-19% lower levels). Interestingly, human Hsp27 extends lifespan when overexpressed in yeast and fly.^30^ Mutation of Hsp27 is associated with the Charcot-Marie-Tooth neuropathy, and Hsp27 plays a role in autophagy and has been recently linked to mitochondrial homeostasis.^31^ In contrast, a few chaperone proteins are also enriched in the insoluble fraction in older mice (Figure S6C-D). Notably, the small heat shock protein CRYAB (a.k.a. HspB5) was the most prominently enriched chaperone protein in the detergent-insoluble fraction from the old cortex and hippocampus. CRYAB has been widely observed to be upregulated or co-aggregated with disease-associated proteins in NDs such as PD, Prion diseases and Alexander’s disease.^32^ This chaperone has also recently been linked to the increased sequestration of insoluble proteins into large cell foci and downregulation of *Cryab* in neurons leads to reduced protein aggregation and increased cytoxocity.^33^ One possibility is that this small heat shock protein has a prominent role in protecting post mitotic cells by sequestering misfolded proteins into insoluble assemblies.

### Feature analysis of datasets with altered PQC

To determine how alterations in PQC might affect the solubility of different groups of proteins, we extended our feature analysis to published datasets using various modulations of chaperones and degradation machinery. Using data of impaired autophagy in mice^34^ and impaired chaperone or proteasome activity in cells^35^, we identified differences between the features associated with proteins that become insoluble due to these stresses. Insoluble proteins resulting from degradation impairment show a greater similarity in feature profiles to the proteins that we identified as insoluble due to aging. Disruption of degradation leads to accumulation of proteins with low intrinsic disorder and higher β-sheet content in the insoluble fraction (Figure 6A). By contrast, inhibition of Hsp70 or Hsp90 generally led to accumulation of proteins with slightly more disorder in the insoluble fraction (Figure 6A). One possibility is that a mild inhibition of these two chaperone systems, potentially caused by the slight reduction in protein levels observed in older mice brain tissues, could lead to the aggregation of a smaller number of clients, possibly more similar to the age-dependent detergent insoluble proteins that we identified. All degradation datasets show depletion for RNA-binding scores consistent with age-related feature analysis (Figure 6A). Together these trends suggest that the accumulation of proteins with these features in the insoluble fraction is most likely due to impaired degradation, possibly of the main targets of PQC pathways.

**Figure 6.**
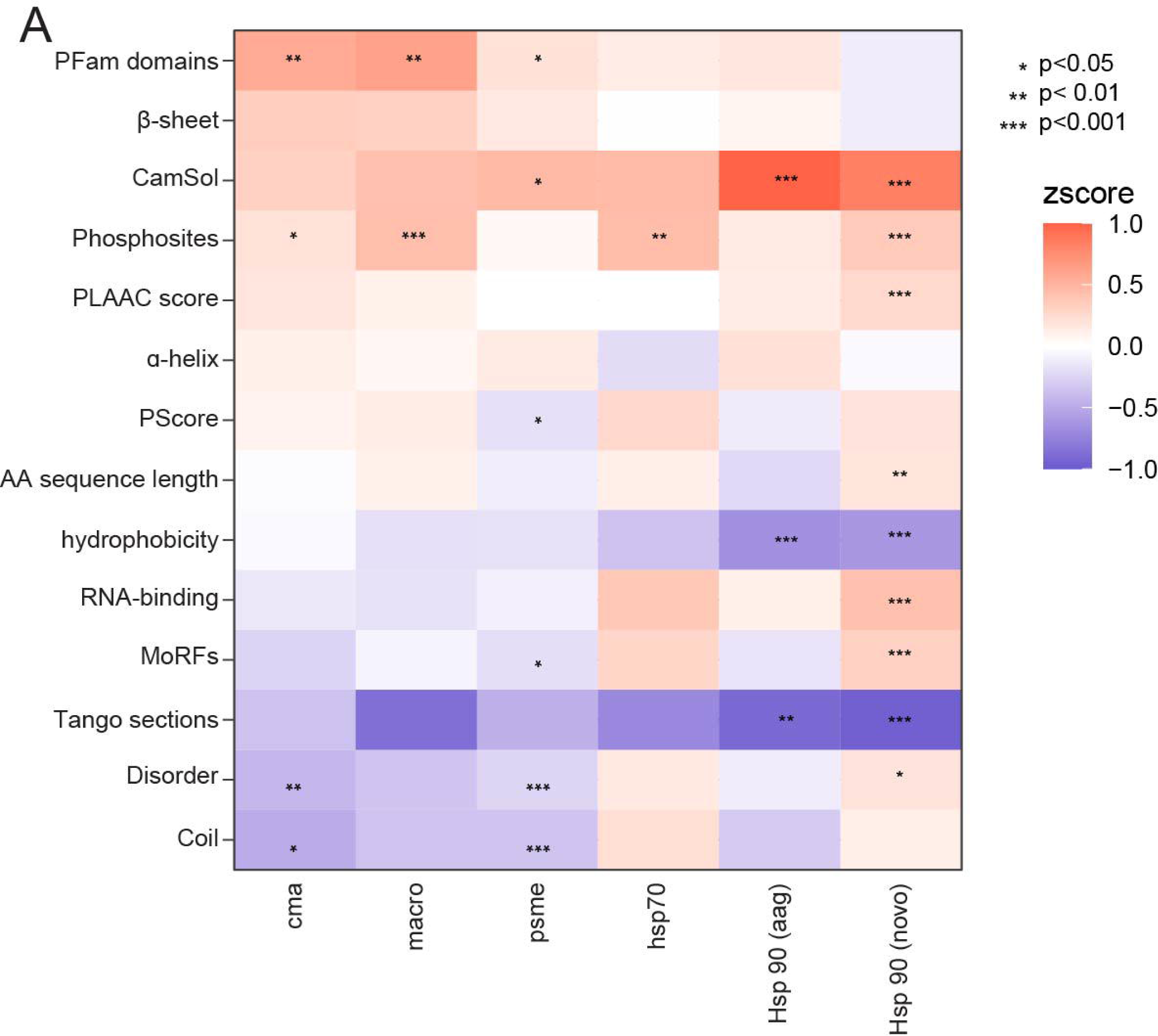
Disruption of protein degradation favors aggregation of proteins with common features. A) Datasets on chaperone mediated autophagy (cma) and macro-autophagy (macro) disruptions (Bourdenx et al), proteasome (psme) inhibition and chaperone inhibition (Sui et al.) were assessed for 14 different parameters using. P-values obtained from Hochberg adjusted Wilcoxon test.

## Discussion

In this study, we compared the composition of proteins enriched in the detergent-insoluble fraction upon aging from two distinct brain regions. While only a subset of proteins was detergent-insoluble in both the cortex and hippocampus, proteins that were insoluble in one brain region typically also trended towards insolubility in the other brain region. Moreover, feature analysis confirmed that proteins that become more insoluble upon aging in each tissue share similar characteristics and are consistent with the detergent-insoluble proteins from our previous analysis using an independent mouse cohort.^12^ Detergent-insoluble protein datasets from young and old mice were distinctly separated by PCA using the same protein features identified in previous studies.^12^ In general, proteins enriched in the detergent-insoluble fraction of older mice were less disordered and had more secondary structure. These findings indicate that these features are consistently associated with proteins that become detergent-insoluble with age across brain regions, potentially due to PQC defects that preferentially effect proteins with more secondary structure.

In order to further investigate changes to proteostasis with age, we assessed the expression and solubility of proteins annotated to specific PQC pathways. Both the cortex and hippocampus have an overall decrease in the expression of chaperone proteins with age. While the overall fold change is low, the reduction of these chaperone levels is significant. One possibility is that a marked decrease of chaperone levels is occurring in only a subset of cells in the older brain tissue, which would then be “masked” by the unaltered chaperone expression of the other cells. In this case, the accumulation of detergent-insoluble proteins may also only occur in a subset of the cells. Alternatively, slightly less chaperones may be required across most cells as a result of the lower population of ribosomes and translation. Interestingly, alterations to ribosome solubility and translation are emerging as a common feature of aging across model organisms.^28^ More work will be required to distinguish between these different possibilities.

Two of the most abundant proteins in the insoluble fraction that are altered upon aging (PLP1 and MOBP) could be components of lipofuscin. Lipofuscin is a mixture of protein and lipids that forms due to incomplete lysosomal degradation.^20^ During aging, microglia are the first cells to accumulate lipofuscin.^16^ Microglia face increased degradation of myelin which can lead to increased lysosomal inclusions during aging.^36^ In additional cell types, incomplete degradation of mitochondria or protein aggregates are additional mechanisms of lipofuscin formation. Impaired degradation leads to accumulation of dysfunctional mitochondria. Accumulation of lipofuscin has been implicated in lysosomal storage disorders and macular degeneration.^37^ Interestingly, lysosomes and mitochondria are two GO terms enriched among the age-dependent detergent-insoluble proteins. Therefore, a subset of observed insoluble proteins could result from incomplete lysosomal degradation, or mislocalization of these proteins in the cytosol.

Both the cortex and hippocampus show strong enrichment of the C1q complex subunits in the detergent insoluble-fraction with age. Complement proteins have frequently been identified as components of aggregates in NDs. Complement may also play a role in age-related decline in memory. C1q has previously been observed to increase in abundance with age in brain tissue and localizes to synapses where it plays a role in synaptic pruning.^22^ In this study we observed an enrichment in the insoluble fraction of C1q components a, b and c. We also observed a decrease in overall synapse protein expression consistent with the role of increased C1q expression in synapse removal. It is possible that these effects are due to the interaction of this complex with antibodies which appear to become insoluble in the brain with age.

Increased inflammation and immunoglobulins have previously been implicated in NDs.^38^ In the data presented here, we find them increased in the aging brain and potentially aggregating in the absence of disease. In MS experiments, several immunoglobulins increased in abundance in older mice and were frequently not measured in young mice. IgM was among the 50 proteins that were enriched in the detergent-insoluble fraction with age in both the cortex and hippocampus. IgGs were also observed to be triton-insoluble by filter trap assay. Microscopy using a separate anti-mouse IgG antibody regularly demonstrated increased background in brain sections from 100-week-old mice while background from 15-week-old mice remained low or undetectable. The IgG signal appears as foci in the brain of old mice. In combination with the filter trap assay these results suggest that immunoglobulins are aggregating in the brains of old mice. However, more analysis would be required to confirm that these foci are aggregates such as co-staining with Thioflavin T or Proteostat. Precedence for the aggregation of antibodies is significant outside of brain tissue. Light chain amyloidosis makes up the largest share of amyloidosis diseases and cases of heavy chain amyloidosis have been reported.^39^ Ig heavy chains form deposits in heavy-chain deposition disease and IgM for non-amyloid Russell bodies in multiple myeloma.^40^ Amyloid plaques that form in early onset AD contain antibodies such as IGKC.^41^ In peripheral tissues, both heavy and light chains are implicated in amyloidosis. Additional experiments are required to confirm the co-aggregation of immunoglobulins and the C1q complex as well as to determine the role that the immunoglobulins might play in the C1q dependent synapse loss observed with age and ND.

## Methods

### Collection of mouse tissues

Male C57BL/6J mice ages 5 (n = 15) and 90 (n = 13) weeks were purchased from the Jackson Laboratories. The authors acknowledge that they did not consider mouse sex differences during experimental design. The mice were held in the UBC Animal Resource Center for an additional 10 weeks under UBC Animal Care Protocol #A18-0067. During this time, they were fed a standard diet (Teklad X2920, Envigo, USA) Mice were sacrificed and processed to obtain whole brain regions and sagittal hemisphere sections as previously described.^12^ Briefly, mice were anesthetized using isoflurane gas followed by and perfusion through the left ventricle with 20mL of 1xPBS containing 1x HALT protease and phosphatase inhibitors. For the lysate-based experiments (MS and FTA), the cortex and hippocampus were collected and snap frozen in liquid nitrogen. Whole left hemispheres were collected for immunofluorescence experiments. These samples were fixed in 4% PFA overnight followed. Subsequently, cryoprotection was done by immersing the tissue in 20% sucrose overnight. Finally, the samples were embedded in OCT media.

### Enrichment for triton X-100 insoluble fraction

Protein extraction from hippocampus and cortex tissue was done on 10 brain samples/day. For each sample, 50 µg of cryoground tissue powder was placed in a 1.5mL lobind tube and brought to 1 mL with lysis buffer (1% triton, 1x PBS, 1x Roche Complete protease inhibitor cocktail, 1 mM PMSF, 2 mM DTT). The tissue was then lysed by bath sonication (Bioruptor, 2×30 seconds on high). The lysates were then centrifuged at 3,000 g for 10 minutes to remove cellular debris. The clarified lysates were removed to a reinforced centrifuge tubes (Beckman Coulter) and centrifuged at 20,000 g for 30 minutes in a Sorvall Legend Micro 21R centrifuge. Concentration measurements were performed by DC protein assay using 2 aliquots from young and old for each fraction. Digestions were done using the entire pellet fraction (∼50 µg) and ∼100 µg of supernatant on S-trap columns.

### Trypsin digests

Protein fractions were reduced with 0.5M DTT and alkylated with 40mM chloroacetamide. Tryptic digests were done on S-trap columns (Protifi). Pellet fractions were resuspended in 5% SDS in 100mM (triethylammonium bicarbonate) TEAB pH 7.5. The entire pellet fraction (∼50 µg) was loaded onto the columns. Supernatant samples (∼50 µg) were combined with an equal volume of 10% SDS in 200mM TEAB pH 7.5. All samples were acidified with phosphoric acid to a final concentration of 1.2%. The samples were combined with S-trap loading buffer (90% MeOH in 100mM TEAB pH 7.1) at a 1:7 ratio and loaded onto the columns. The columns were washed 3x with S-trap loading buffer, 1x with MeOH/CHCl_3_ and then 2 additional times with S-trap loading buffer. Trypsin (Promega) in 50mM TEAB pH 7.1 was added at a 1:25 ratio. The digestion buffer was centrifuged into the column and the reaction was incubated overnight at 37°C. The resulting peptides were eluted from the column and acidified with 10% TFA to pH <3. Peptides were then desalted using C18 stage tips.^42^

### MS data acquisition and data analysis

MS analysis was performed as previously described.^12^ Detergent solubility samples were run on a Q exactive HF mass spectrometer. Online fractionation was performed with an Easy-nLC 1200 liquid chromatography system (Thermo Scientific) configured with a 20µl sample loop, 30 µm ID steel emitter and a 50 cm µPACTM analytical column (Pharmafluidics) with trap column heated to 50°C. The LC gradient ranged from 4-27% buffer B (95% ACN in 0.1% FA in water) in combination with buffer A (2% Acetonitrile (ACN) in 0.1% FA in water). For each sample 800ng were injected. Data files were searched using Spectronaut (version 17.6.230428.55965) in Direct DIA mode. A mouse FASTA file (Swiss-Prot reviewed, not containing isoforms) was used as the sequence database (January 3^rd^, 2023). Methionine oxidation and protein N-term acetylation were included as variable modifications. Carbamidomethyl cysteine was set as a fixed modification.

### Immunofluorescence in tissue sections

12 µm left hemisphere brain sections were generated by cryosectioning (CM 3050C cryostat, Leica, Germany). Sections were thaw-mounted on ProbeOn Plus charged slides (Fisher, USA). Slides containing brain sections were dried overnight at room temperature. The slides were then washed 2x for 5 minutes each in 1xTBS-T. Antigen retrieval was done by incubating the slides in 10 mM Sodium citrate pH 6 with 0.05% Tween-20 in boiling water for 30 minutes in a pressure cooker. The sections were then blocked with 5% goat serum and permeabilized with 0.1% triton in 1x PBS for 1 hour at RT. Secondary antibody staining was done with a polyclonal goat anti-mouse IgG conjugated to Alexa 568 (Invitrogen A11011; 1:200). The secondary antibody was incubated for 90 minutes at 4°C followed by washing three times for 5 minutes each. The sections were then incubated with DAPI for 10 minutes. Finally, the slides were mounted with Dako fluorescent mounting media (Molecular Probe, Eugene, OR). All incubations were done in a humidified slide box. Imaging of the cortex was done with an Olympus FV1000 microscope using a 10x objective (UPLSAPO60XW).

### Filter trap assay

We performed the FTA as previously described.^12^ Briefly, mouse cortex tissue lysates (triton X-100) were centrifuged at 3,000 RCF for 10 minutes and the resulting supernatants were diluted to 0.4µg/µL. Dilutions were done with either triton lysis buffer or with SDS lysis buffer for a final concentration of 1% triton or 2% SDS. 100µL (2 × 50µL) were passed through a 0.2µm cellulose acetate membrane and Whatman filter paper fitted in a Scie-Plas dotblot apparatus with vacuum. Wells were washed 3x with 50µL of the corresponding buffer and membranes were blocked for 1 hour in 5% milk in TBS-T followed. Goat anti-mouse IgG (LiCor 926-32210) was diluted 1:10,000 and incubated with the membrane for 1 hour at room temperature. Imaging was performed with the Odyssey system (LiCor), Quantification was done with the Image Studio (LiCor).

### Computational analysis

GO analysis of significantly enriched hits was performed using DAVID 2021 with the default parameters. Clustering of similar GO terms was done by DAVID and representative terms were chosen from clusters with a fold enrichment greater than two were and FDR corrected p-values less than 0.05. Additional analysis of terms related to PQC was done using downloaded Quick GO annotations. Specific chaperone classes were annotated by Shemesh et al.^43^ The human chaperone UniProt codes were converted to mouse using Mouse Genome Informatics homologies (Jackson Laboratory). The feature analysis was handled using an in-house R script established for a recent study.^12^ Significance was determined using either a Wilcoxon test or a Fischer test. P-values were corrected using the p.adjust function in R applying an FDR (Benjamini-Hochberg) correction. Protein solubility scores calculated were obtained from CamSol.^44^ Intrinsic protein disorder was calculated with DISOPRED3.^45^ Hydrophobicity calculations were obtained from the grand average hydrophobicity score (GRAVY).^46^ Scores RNA-binding likelihood were calculated by RBPPRED ^47^.Calculations of Protein secondary structure were performed using SCRATCH.^48^ Prediction of aggregation-prone regions was obtained from TANGO scores.^49^ Abundances were gathered from PaxDB and log transformed.^50^ Pfam domain annotations (version 33.0) were downloaded and processed in R to calculate the percentage of each sequence in Pfam domains.^51^ Annotated phosphorylation site information was obtained from PhosphoSite Plus.^52^ R scripts used to analyze data can be found at https://github.com/cmolzahn/aging_proteomics.

## Abbreviations

AD: Alzheimer’s disease
DIA: data-independent acquisition
FTA: filter trap assay
GO: gene ontology
MLO: membraneless organelle
MS: mass spectrometry
ND: neurodegenerative disease
NFTs: neurofibrillary tangles
PCA: principal component analysis
PQC: protein quality control
TEAB: triethylammonium bicarbonate

## Acknowledgments

We thank Irina Zemlyankina and Drs. Tahir Ali and Sabine Gilch for support preparing cryosections, Samuel Leung and Dr. Gabriela Cohen Freue for data analysis discussion, members of the Mayor lab for discussions, the UBC bioimaging facility and the proteomics facility for help and support.

---

**Figure S1.**
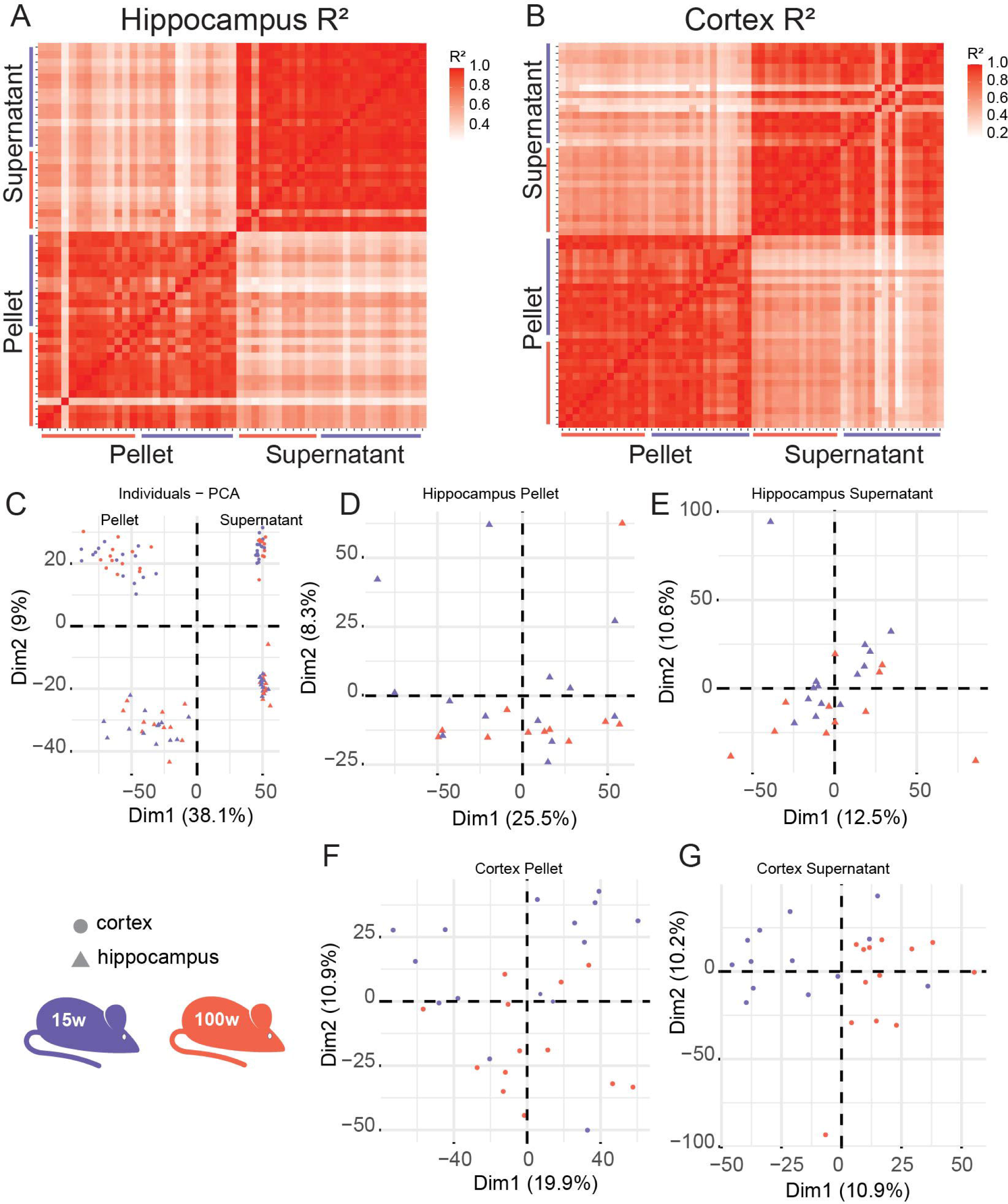
Proteins show strong clustering by solubility and brain region. Heatmaps of R^2^ values show strong correlation between replicates within solubility groups in the A) Hippocampus and B) Cortex. C) PCA applied to all data shows strong clustering by both the solubility fraction and the brain region. D-G) PCA applied within only one brain region and fraction confirms that the samples do not cluster by age in the pellet fractions (D, F) or supernatant fractions (E, G).

**Figure S2.**
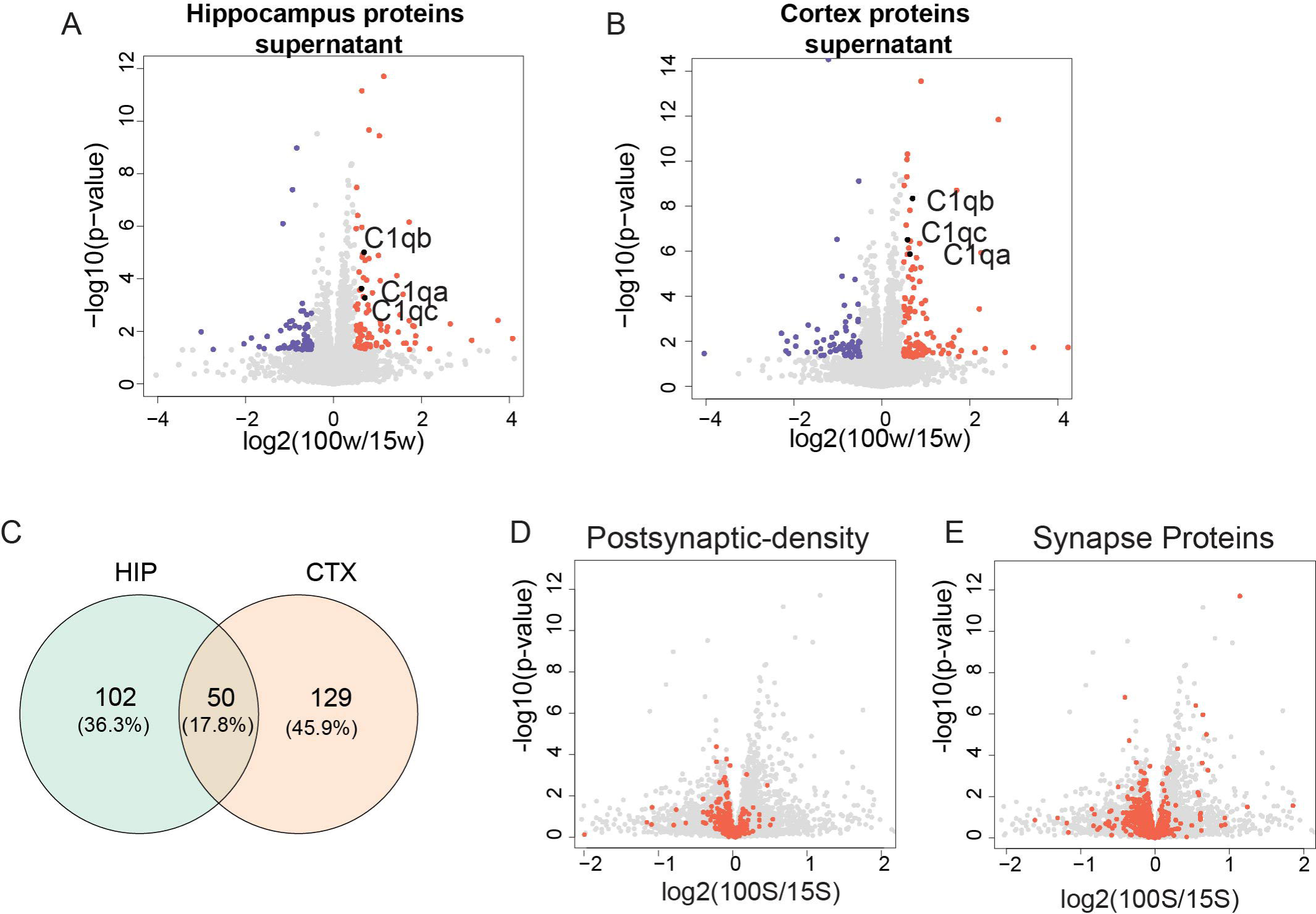
Altered soluble proteins identified in both the cortex and hippocampus. Significantly altered proteins between mice ages 100- and 15-weeks-old identified in the A) hippocampus supernatant fraction and B) cortex supernatant fraction. Significance was determined using a two-sample t-test assuming equal variance. The p-value cut off was set as 0.05 and the fold change at a log_2_ ratio of 0.5. Complement C1q complex proteins are highlighted. C) Venn diagram representing the proteins identified as enriched in the detergent-insoluble fraction with age in both brain regions tested. D-E) Abundance of post-synaptic density (D) and synapse proteins (E) in hippocampus tissues are highlighted in red.

**Figure S3.**
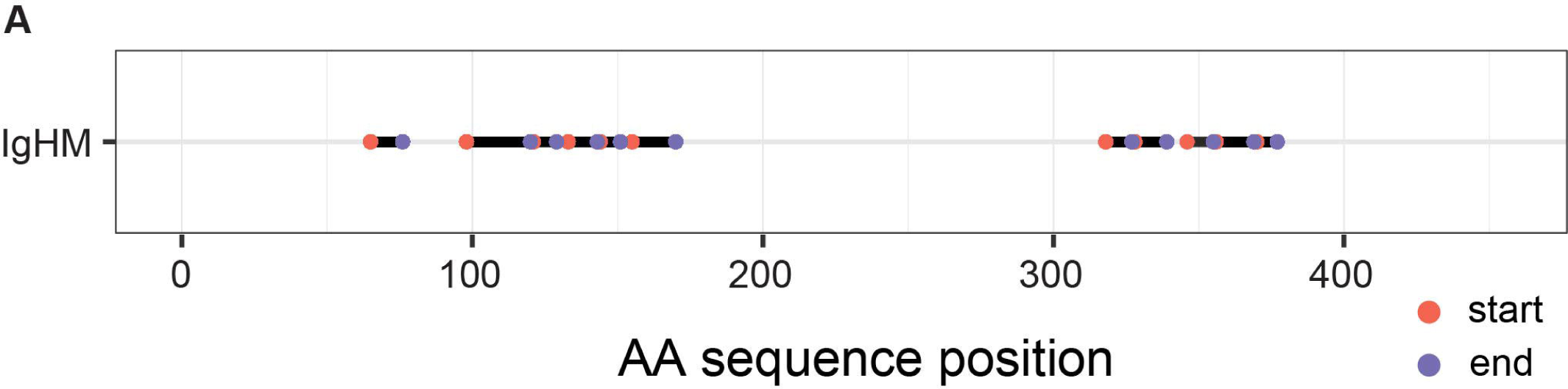
Position of IgHM peptides along the amino acid sequence. All identified peptides that were mapped to IgHM are shown. The start position of each peptide is shown in red and the end position is shown in purple.

**Figure S4.**
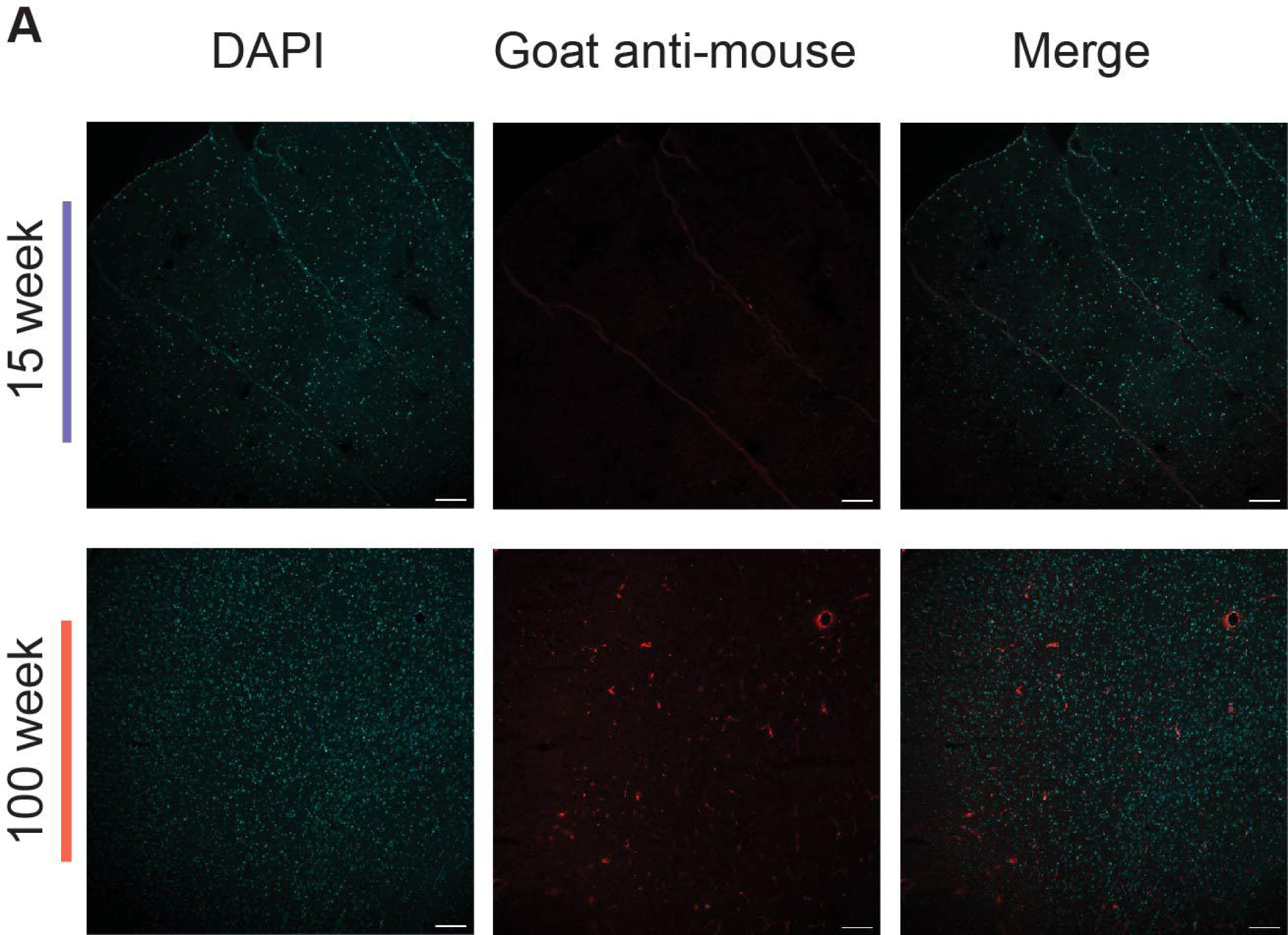
Increased immunoglobulin signal observed in the aging mouse cortex. Mouse cortex stained with goat anti-mouse IgG (1:200) and DAPI taken with a 10x objective from a young mouse (mo 11) and an old mouse (mo 28). Scale bar is 200µm.

**Figure S5.**
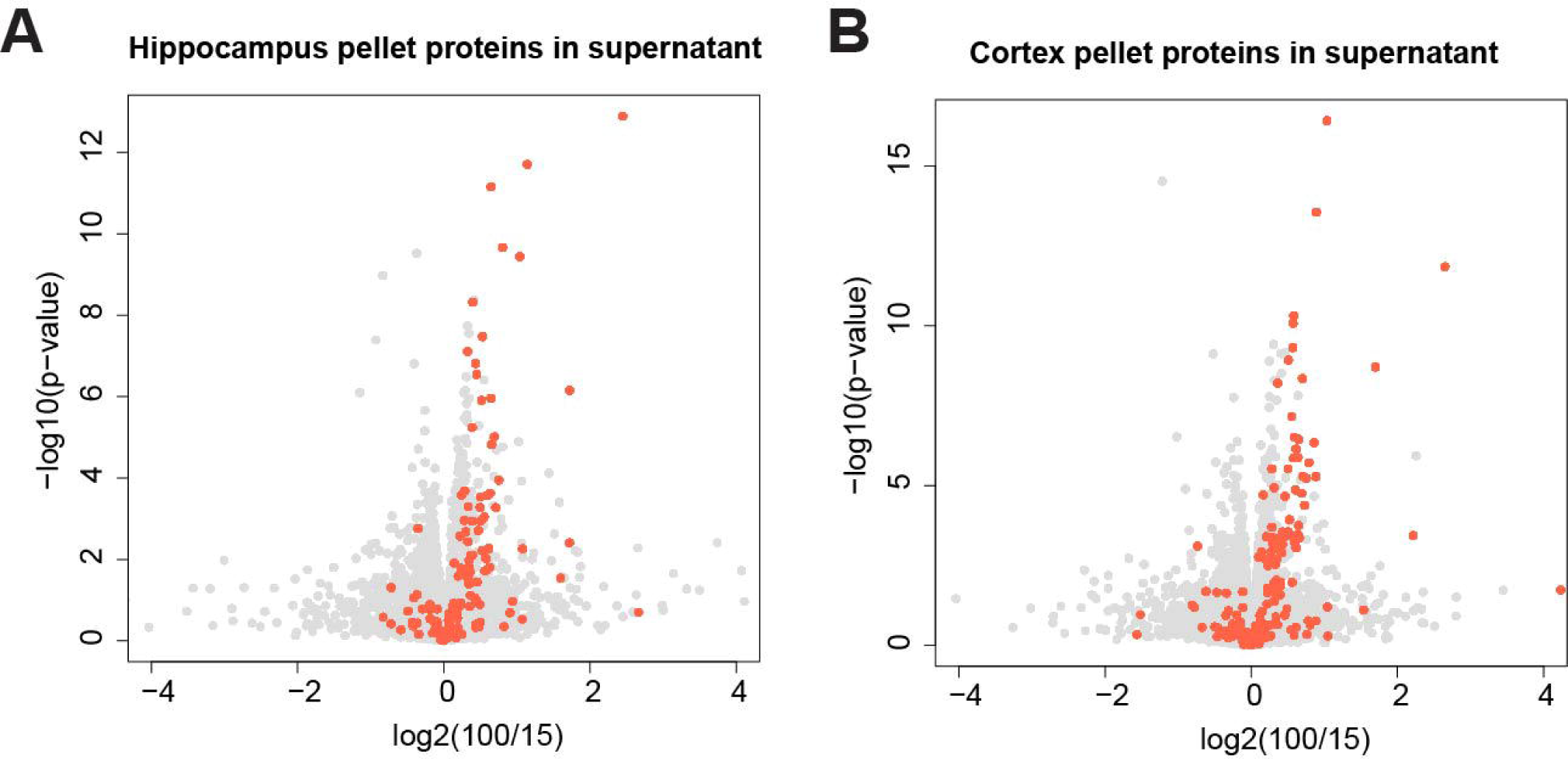
Insoluble proteins tend to be more abundant. Detergent-insoluble proteins are mapped to their corresponding values in the supernatant in the A) Hippocampus and B) cortex. P-values obtained from a two-sample unpaired t-test.

**Figure S6.**
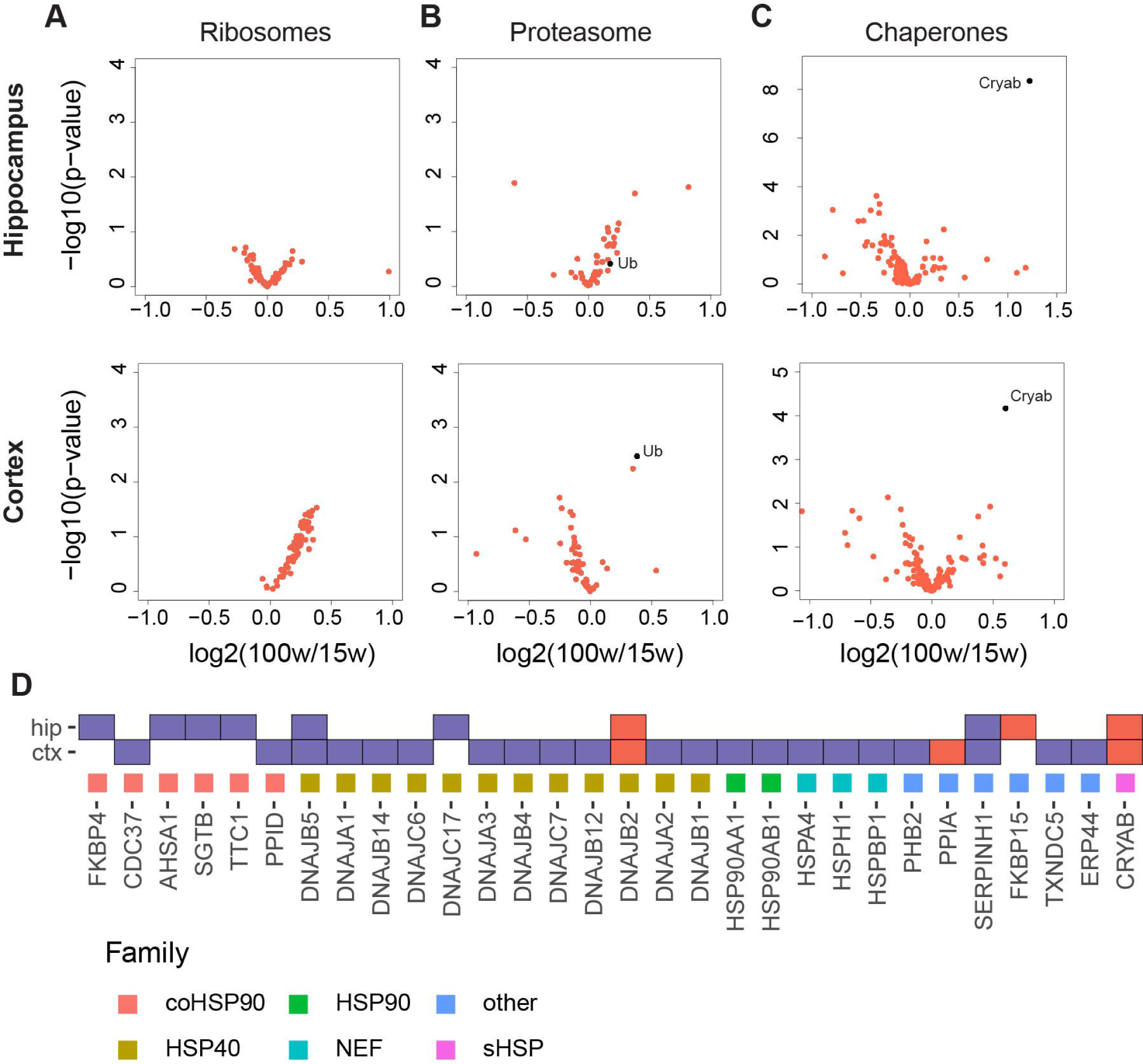
Age-associated alterations in the solubility of PQC components. A-C) Proteins annotated with the indicated GO terms for in each sample were plotted according to their log_2_(fold-change) and t-test-derived p-value to highlight changes in their levels in the pellet fractions for the hippocampus (top row) and the cortex (bottom row) ribosomal proteins (A), proteasomal subunits and ubiquitin (Ub) (B), and chaperones (C). D) Chaperone significantly depleted (purple) or enriched (red) (p-value < 0.05) in the older hippocampus (HIP) and cortex (CTX).

